## Supplemental Materials for "Evaluating how lethal management affects poaching of Mexican wolves"

^2^ Project Coyote Science Advisory Board

^*^Equal first co-authors

Diagnostic step: Analysis of Aggregated Human-Caused Endpoint

Here, we report the results of the Cox model and FG competing risk model for the aggregated human-caused mortality endpoint. Periods of liberalized wolf-killing were associated with a 1% increase in hazard (HR=1.01) for the aggregated human-caused mortality endpoint, relative to periods of stricter protection, compatible with a range that overlaps zero of -44% to +56% (p=0.935). The proportion of collared wolves (CIF) with the aggregated human-caused endpoint decreased by 12% (SHR= 0.88, compatible interval=-44% to +38%, p=0.579). Therefore, our presumption that disaggregating the human causes of mortality by criminal and non-criminal endpoints would prove more informative seems justified.

Supplementary Figs. S1 – S7

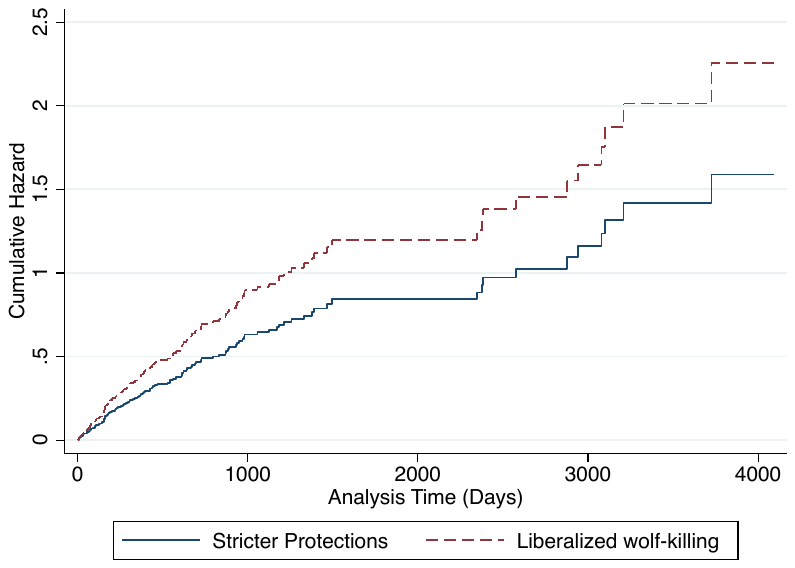

**Supplementary Fig. S1.** Cumulative hazard function for the total potential poached (LTF + reported poached) endpoint for n = 119 collared Mexican wolves. Lines show cumulative hazard over monitoring time derived from a univariate Cox model for 2 policy periods of baseline strict protections (solid blue line) relative to intervention or liberalized wolf-killing periods (dashed red line) with an HR=1.42 over the baseline.

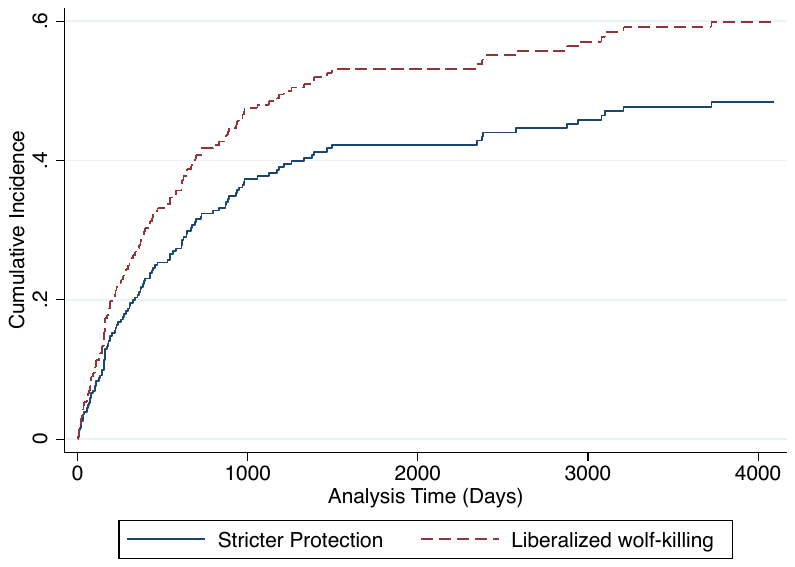

**Supplementary Fig. S2.** Cumulative incidence function for the total potential poached endpoint (LTF + reported Poached) for n=119 collared Mexican wolves. Curves show incidence over monitoring time derived from a Fine-Grey subhazard model for 2 policy periods of baseline strict protection (solid blue line) relative to intervention or liberalized killing periods (dashed red line) with an SHR = 1.38 over the baseline.

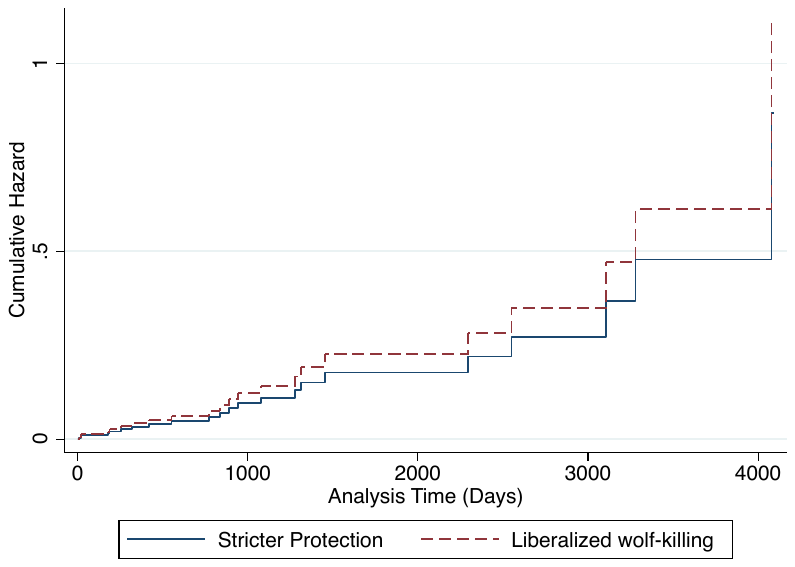

**Supplementary Fig. S3.** Cumulative hazard function for the natural endpoint for n = 22 collared Mexican wolves. Lines show cumulative hazard over monitoring time derived from a univariate Cox model for 2 policy periods of baseline strict protections (solid blue line) relative to intervention or liberalized killing periods (dashed red line) with an HR=1.28 over the baseline.

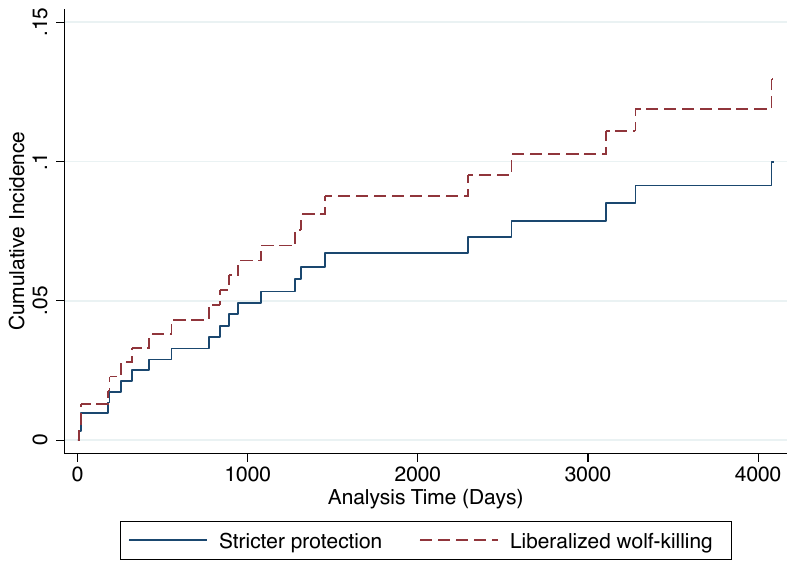

**Supplementary Fig. S4.** Cumulative incidence function for the natural endpoint for n=22 collared Mexican wolves. Curves show incidence over monitoring time derived from a Fine-Grey subhazard model for 2 policy periods of baseline strict protection (solid blue line) relative to intervention or liberalized killing periods (dashed red line) with an SHR = 1.32 over the baseline.

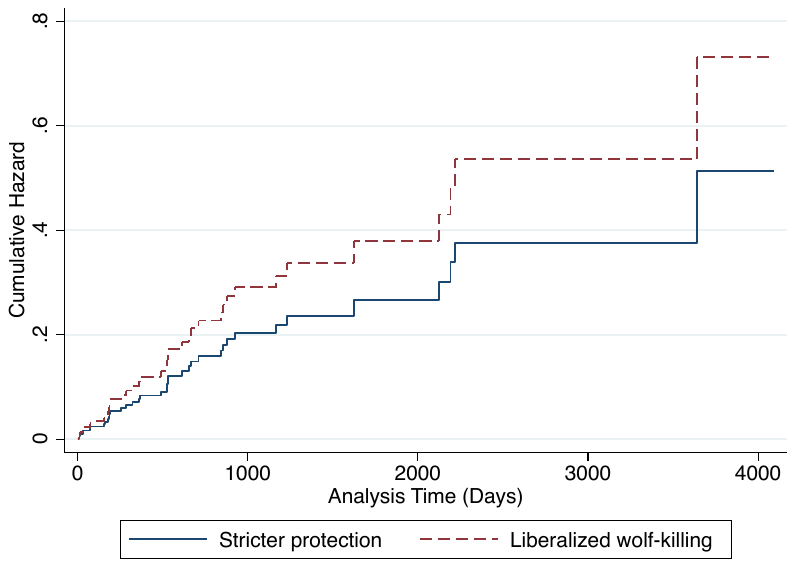

**Supplementary Fig. S5.** Cumulative hazard function for the non-criminal endpoint for n = 38 collared Mexican wolves. Lines show cumulative hazard over monitoring time derived from a univariate Cox model for 2 policy periods of baseline strict protections (solid blue line) relative to intervention or liberalized killing periods (dashed red line) with an HR=1.42 over the baseline.

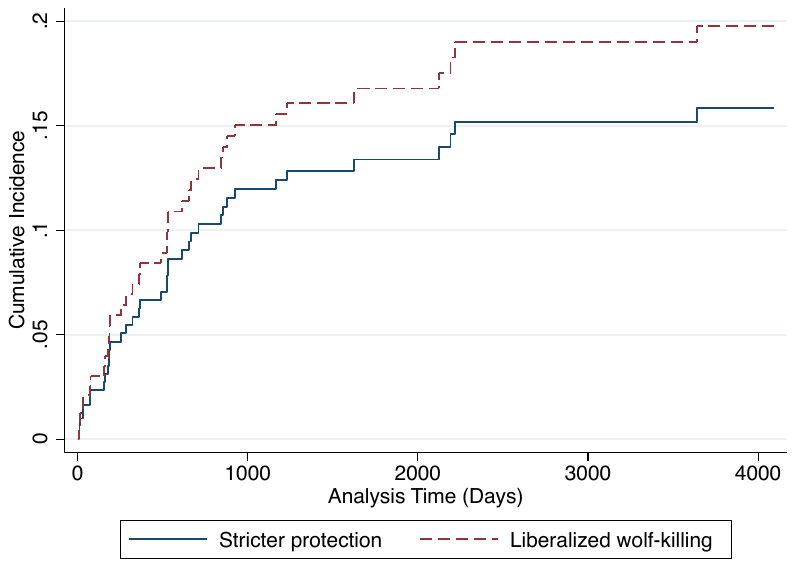

**Supplementary Fig. S6.** Cumulative incidence function for the non-criminal endpoint for n=38 collared Mexican wolves. Curves show incidence over monitoring time derived from a Fine-Grey subhazard model for 2 policy periods of baseline strict protection (solid blue line) relative to intervention or liberalized killing periods (dashed red line) with an SHR = 1.27 over the baseline.

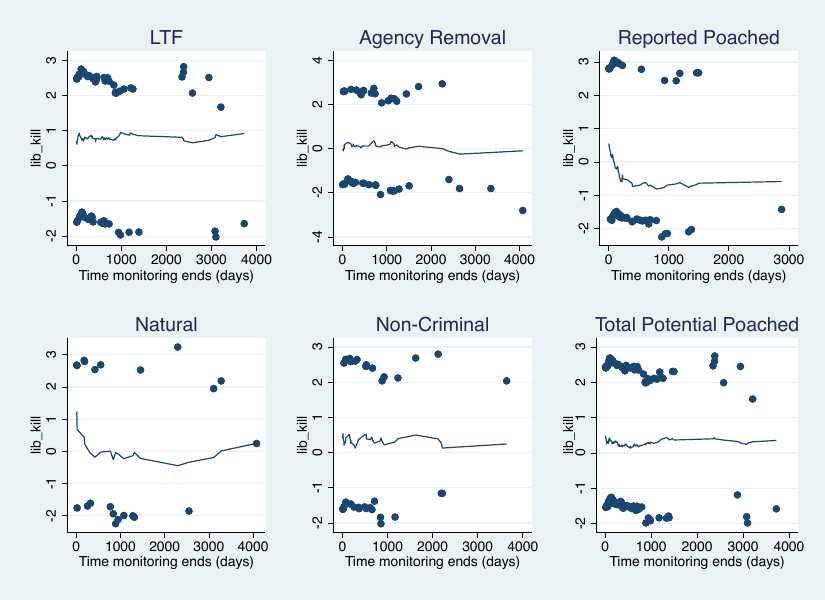

**Supplementary Fig. S7.** Scaled Schoenfeld residual scatterplot for each endpoint in the best Cox model. A line with slope of zero between the points suggests the log hazard-ratio is proportional over time and therefore that the proportional hazard assumption is met.

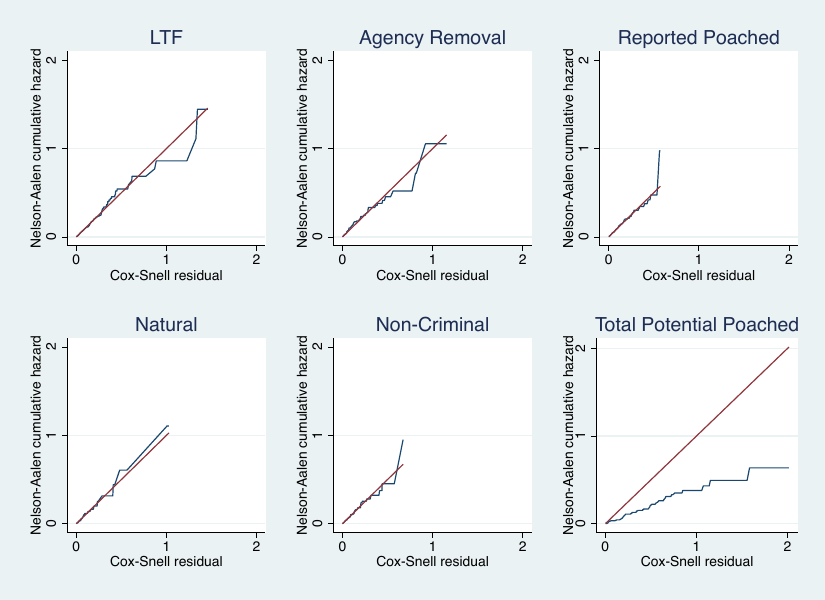

**Supplementary Fig. S8.** Cox-Snell generalized residuals (red lines) are used for evaluating the goodness of fit of the best Cox model to the individual wolf survival data. Blue lines represent the Nelson-Aalen cumulative hazard. Good model fit is suggested when the blue and red lines overlap closely. Total potential poached had the model with the worst fit, likely because LTF and reported poached hazard models had opposite predicted directions.

Supplementary Tables S1-S2

**Supplementary Table S1.** Model parameters for our policy intervention (*lib_kill*) from Cox and Fine-Grey models for 279 collared, adult Mexican wolves. Comp. Int refers to the compatible interval around point estimates. The best model is marked in bold-faced text.

| **Model** | **Cox Model** | | | | | **Fine-Grey Model** | | | | |
| --- | --- | --- | --- | --- | --- | --- | --- | --- | --- | --- |
|  | **HR** | **Comp. Int.** | **p-value** | **AIC** | **BIC** | **SHR** | **Comp. Int.** | **p-value** | **AIC** | **BIC** |
|  | **LTF** | | | | | | | | | |
| **Liberalized Killing Periods (lib_kill)** | **2.21** | **1.36-3.60** | **0.001** | **597.73** | **603.08** | **2.28** | **1.38-3.76** | **0.001** | **691.28** | **696.63** |
| +Season (winter) | 2.15 | 1.32-3.51 | 0.002 | 598.18 | 608.88 | 2.24 | 1.35-3.72 | 0.002 | 692.52 | 703.22 |
| +Sex (female) | 2.18 | 1.34-3.54 | 0.002 | 597.69 | 608.40 | 2.31 | 1.40-3.81 | 0.001 | 689.93 | 700.64 |
|  | **Agency Removal** | | | | | | | | | |
| **Liberalized Killing Periods (lib_kill)** | **1.05** | **0.588-1.88** | **0.86** | **426.33** | **431.68** | **0.96** | **0.525-1.75** | **0.892** | **506.59** | **511.95** |
| +Season (winter) | 1.07 | 0.599-1.91 | 0.818 | 427.68 | 438.38 | 0.98 | 0.538-1.79 | 0.953 | 507.52 | 518.22 |
| +Sex (female) | 1.05 | 0.589-1.88 | 0.86 | 428.33 | 439.04 | 0.96 | 0.523-1.75 | 0.888 | 508.58 | 519.28 |
|  | **Reported Poached** | | | | | | | | | |
| **Liberalized Killing Periods (lib_kill)** | **0.78** | **0.437-1.39** | **0.407** | **505.61** | **510.96** | **0.69** | **0.378-1.25** | **0.226** | **554.27** | **559.61** |
| +Season (winter) | 0.77 | 0.434-1.38 | 0.389 | 507.30 | 518.00 | 0.69 | 0.376-1.25 | 0.219 | 556.21 | 566.90 |
| +Sex (female) | 0.78 | 0.437-1.39 | 0.400 | 506.49 | 517.19 | 0.68 | 0.373-1.23 | 0.204 | 554.75 | 565.45 |
|  | **Total Potential Poached (LTF + Reported Poached)** | | | | | | | | | |
| **Liberalized Killing Periods (lib_kill)** | **1.42** | **0.993-2.03** | **0.055** | **1108.81** | **1114.16** | **1.38** | **0.946-2.01** | **0.095** | **1215.10** | **1220.45** |
| +Season (winter) | 1.39 | 0.973-1.99 | 0.070 | 1109.03 | 1119.73 | 1.35 | 0.927-1.99 | 0.116 | 1216.24 | 1226.95 |
| +Sex (female) | 1.41 | 0.991-2.03 | 0.056 | 1110.62 | 1121.32 | 1.38 | 0.948-2.02 | 0.092 | 1216.82 | 1227.52 |
|  | **Non-Criminal** | | | | | | | | | |
| **Liberalized Killing Periods (lib_kill)** | **1.42** | **0.750-2.71** | **0.278** | **353.11** | **358.46** | **1.28** | **0.663-2.46** | **0.463** | **405.04** | **410.39** |
| +Season (winter) | 1.37 | 0.724-2.61 | 0.331 | 353.27 | 363.97 | 1.24 | 0.643-2.39 | 0.521 | 405.76 | 416.46 |
| +Sex (female) | 1.43 | 0.752-2.74 | 0.274 | 353.98 | 364.69 | 1.27 | 0.660-2.43 | 0.476 | 405.99 | 416.69 |
|  | **Natural** | | | | | | | | | |
| **Liberalized Killing Periods (lib_kill)** | **1.28** | **0.531-3.08** | **0.582** | **181.53** | **186.88** | **1.32** | **0.559-3.12** | **0.526** | **232.70** | **238.05** |
| +Season (winter) | 1.27 | 0.522-3.07 | 0.60 | 182.57 | 193.28 | 1.29 | 0.528-3.13 | 0.579 | 234.24 | 244.94 |
| +Sex (female) | 1.28 | 0.533-3.06 | 0.583 | 183.52 | 194.22 | 1.32 | 0.553-3.15 | 0.531 | 234.70 | 245.41 |

**Supplementary Table S2.** parameters for the Bayes Factor Calculator under each of the three specifications and for each endpoint relevant to hypotheses [46]. Standard error (SE) values are calculated by dividing the sample mean by the z-score of the model coefficient. Sample means are ln(HR) or ln(SHR) for each endpoint from Tables 5 and 6 respectively. Theory mean and theory standard deviation (SD) refer to the predicted HR and SHRs used to create the half normal likelihood functions under specifications (1) and (3) in Methods. Theory SD and Upper bounds are ln(HR) or ln(SHR) from tables 5 and 6 respectively. Specification (3) uses analogous estimates of HR and SHR from Santiago-Avila et al. [18]. Standard deviations and upper bounds cannot be negative values, therefore we took the absolute value when necessary. MX = Mexican wolf, WI = Wisconsin grey wolf in reference to [18]. POA = reported poached. LTF+POA = total potential poached

| **BF Specifications** | **HR** | | | | | | **SHR** | | | | | |
| --- | --- | --- | --- | --- | --- | --- | --- | --- | --- | --- | --- | --- |
|  | **Inputs** | | | | | **BF** | **Inputs** | | | | | **BF** |
|  | **SE** | **Sample Mean** | **Lower-Upper Bound** | **Theory Mean** | **Theory SD** |  | **SE** | **Sample Mean** | **Lower-Upper Bound** | **Theory Mean** | **Theory SD** |  |
|  | **LTF** | | | | | | | | | | | |
| (1) half-normal w/MX-agency removal | 0.247 | 0.797 | - | 0 | 0.051 | 1.80 | 0.259 | -0.838* | - | 0 | \|-0.042\| | 0.69 |
| (2) uniform w/upbound-MX agency removal | 0.247 | 0.797 | 0-0.051 | - | - | 1.41 | 0.259 | 0.838 | 0-\|-0.042\| | - | - | 1.30 |
| (3) half-normal w/WI LTF | 0.247 | 0.797 | - | 0 | 0.166 | 8.08 | 0.259 | 0.838 | - | 0 | 0.174 | 8.08 |
| **BF Specifications** | **Reported Poached** | | | | | | | | | | | |
| (1) half-normal w/MX-agency removal | 0.296 | -0.246 | - | 0 | 0.051 | 0.89 | 0.307 | \|-0.371\|** | - | 0 | \|-0.042\| | 1.14 |
| (2) uniform w/upbound-MX agency removal | 0.296 | -0.246 | 0-0.051 | - | - | 0.93 | 0.307 | -0.371 | 0- \|-0.042\| | - | - | 0.92 |
| (3) half-normal w/WI POA | 0.296 | \|-0.246\|** | - | 0 | \|-0.210\| | 1.25 | 0.307 | \|-0.371\|** | - | 0 | \|-0.274\| | 1.63 |
| **BF Specifications** | **Total Potential Poached (LTF + Reported Poached)** | | | | | | | | | | | |
| (1) half-normal w/MX-agency removal | 0.313 | 0.350 | - | 0 |  | 1.15 | 0.193 | -0.322* | - | 0 | \|-0.042\| | 0.76 |
| (2) uniform w/upbound-MX agency removal | 0.313 | 0.350 | 0-0.051 | - | - | 1.09 | 0.193 | 0.322 | 0- \|-0.042\| | - | - | 1.19 |
| (3) half-normal w/WI LTF+POA | 0.313 | 0.350 | - | 0 | 0.131 | 1.35 | 0.193 | 0.322 | - | 0 | 0.122 | 2.05 |

* For positive sample means paired with negative predicted means, we changed the sign of the sample mean.

**For negative sample means paired with negative predicted mean, we took the absolute value of both variables to represent the same effect direction.

Statistical code (STATA)

Statistical code for all analyses conducted in STATA.

**Mexican Wolf Survival Analysis**

***Data Setup***

********************************************************************************

***DATA EXPLORATION & SETUP*****************************************************

********************************************************************************

gen data_order = _n

order data_order, before(wolf_ID)

bysort wolf_ID: gen nvals = _n == 1

count if nvals

/*NOTES -- wolf_ID 1289/1295/1340 are capture deaths; 1438 is a pup and was dropped*/

**VARIABLES NEEDED (for setup)***

*Origin of full monitoring history for all spells --> monit_origin_date

by wolf_ID, sort: egen monit_origin_date = min(spell_start_date)

order monit_origin_date, after(sex)

*switch var format to DATE*

format %tdnn/dd/CCYY monit_origin_date

*End of full monitoring history for all spells --> date_endpoint

by wolf_ID, sort: egen date_endpoint = max(spell_end_date)

order date_endpoint, after(spell_end_date)

*switch var format to DATE*

format %tdnn/dd/CCYY date_endpoint

*Sex

gen sex_coded = 0

replace sex_coded = 1 if sex == "f"

tab sex_coded sex

order sex_coded, after(sex)

*Generate identifier for spell ending

gen cause_spell_end = "agency removal" if removed==1

replace cause_spell_end = "missing" if missing==1

replace cause_spell_end = "natural death" if natural_mort==1

replace cause_spell_end = "human-caused death" if mortality==1 & OK_hum_caused==1

*Generate aggregate cause_endpoint variable with JO Survival data

gen cause_endpoint_JO = cause_spell_end if spell_end_date==date_endpoint

replace cause_endpoint_JO = "LTF" if cause_endpoint_JO=="missing"

replace cause_endpoint_JO = "CENSORED" if spell_end_date>=20819

**Encode cause_endpoint variable from JO and merge with OLE data

encode cause_endpoint_JO, gen(cause_endpoint_JO_coded)

tab cause_endpoint_JO_coded

gen month_endpoint = month(spell_end_date) if cause_endpoint_JO_coded!=.

order month_endpoint, after(date_endpoint)

tab cause_endpoint_JO_coded month_endpoint if year(date_endpoint)==2005, mi

tab cause_endpoint_JO_coded month_endpoint if year(date_endpoint)==2006, mi

tab cause_endpoint_JO_coded month_endpoint if year(date_endpoint)==2007, mi

tab cause_endpoint_JO_coded month_endpoint if year(date_endpoint)==2008, mi

tab cause_endpoint_JO_coded month_endpoint if year(date_endpoint)==2009, mi

merge m:1 wolf_ID using "Cause_of_death_OLE.dta", gen(OLE_merge)

list wolf_ID if OLE_merge==2

drop if OLE_merge==2

order cause_endpoint_OLE, after(cause_endpoint_JO_coded)

order OLE_merge, after(cause_endpoint_OLE)

replace cause_endpoint_OLE="" if cause_endpoint_JO_coded==.

tab cause_endpoint_OLE

tab cause_endpoint_OLE cause_endpoint_JO_coded, mi

gen cause_endpoint_disag = cause_endpoint_JO if cause_endpoint_JO_coded!=.

replace cause_endpoint_disag = "illegal take" if cause_endpoint_disag=="human-caused death" & cause_endpoint_OLE=="Gunshot" | cause_endpoint_disag=="human-caused death" & cause_endpoint_OLE=="Gunshot/Blunt force trauma" ///

| cause_endpoint_disag=="human-caused death" & cause_endpoint_OLE=="Trap injuries" | cause_endpoint_disag=="human-caused death" & cause_endpoint_OLE=="Blunt force trauma" | cause_endpoint_disag=="human-caused death" & cause_endpoint_OLE=="Unknown" & wolf_ID==647 | cause_endpoint_disag=="human-caused death" & cause_endpoint_OLE=="Unknown" & wolf_ID==756

*2 Unknowns --> 'illegal' wolf 647 never recovered (collar cut off); 756 too decomposed for necropsy

*Remaining human-caused deaths classified as as non-criminal

replace cause_endpoint_disag = "non-criminal" if cause_endpoint_disag=="human-caused death"

order cause_endpoint_disag, after(OLE_merge)

*final cause_endpoint var

tab cause_endpoint_disag

encode cause_endpoint_disag, gen(cause_endpoint_enc)

order cause_endpoint_enc, after(cause_endpoint_disag)

tab cause_endpoint_enc cause_endpoint_disag

********************************************************************************

*****TIME-SPLITTING DATASET FOR COMPETING RISK ANALYSES*************************

********************************************************************************

*replace date_endpoint=. if cause_endpoint_enc==.

*replace monit_origin_date=. if monit_origin_date!=spell_start_date

stset spell_end_date, failure(cause_endpoint_enc==2 3 4 5 6) exit(failure) origin(time monit_origin_date) time0(spell_start_date) id(wolf_ID)

list wolf_ID _t0 _t _d _st in 1/20

stdes

stsum

*****TIME-SPLITTING for time-dependent variables and 'spells'*****

*****TREATMENT VARIABLE*****

*10/10/2005 - 16719

*11/13/2009 - 18233

stsplit treat_split, at(16719 18233 20104) after(monit_origin_date==1/1/1960)

***Generating liberalized killing treatment binary variable (1 if lib kill period)

gen lib_kill = 0

replace lib_kill = 1 if treat_split==16719 | treat_split==20104

gen treat_periods = 1

replace treat_periods = 2 if spell_start_date>=16719

replace treat_periods = 3 if spell_start_date>=18233

replace treat_periods = 4 if spell_start_date>=20104

*Reg report Table 2

tab cause_endpoint_JO_coded treat_period

tab lib_kill if _d==1

tab cause_endpoint_JO_coded lib_kill

*Reg report Table 3

tab cause_endpoint_enc treat_period

tab cause_endpoint_enc lib_kill

*aggregate policy period summary stats

bysort lib_kill: stsum

bysort treat_period: stsum

*****WINTER-SUMMER VARIABLE*****

stsplit season_split, at(13969	14152	14334	14517	14700	14883	15065	15248	15430	15613	15795	15978	16161	16344	16526	16709	16891	17074	17256	17439	17622	17805	17987	18170	18352	18535	18717	18900	19083	19266	19448	19631	19813	19996	20178	20361	20544	20727) after(monit_origin_date==1/1/1960)

**Generating winter (1) time-dep binary variable

gen winter = 0

replace winter = 1 if season_split==14152 | season_split==14517 | season_split==14883 | ///

season_split==15248 | season_split==15613 | season_split==15978 | season_split==17805 | ///

season_split==16709 | season_split==17074 | season_split==17439 | season_split==17837 | ///

season_split==18170 | season_split==18535 | season_split==18900 | season_split==19266 | ///

season_split==19631 | season_split==19996 | season_split==20361 | season_split==20727

tab winter if _d==1

tab lib_kill winter

tab lib_kill winter if _d==1

tab cause_endpoint_JO_coded winter if _d==1

tab cause_endpoint_enc winter if _d==1

***Survival Analysis***

********************************************************************************

***CHECKING FOR CAUSE-SPECIFIC DIFFERENCES IN HAZARD & PH ASSUMPTION************

********************************************************************************

**ALL EVENTS**

stset spell_end_date, failure(cause_endpoint_enc==2 3 4 5 6) exit(failure) origin(time monit_origin_date) time0(spell_start_date) id(wolf_ID)

stdes

stsum

*logrank test

sts test lib_kill, strata(cause_endpoint_enc) d

********************************************************************************

****ST COX FOR CAUSE-SPECIFIC HAZARDS*******************************************

********************************************************************************

***LTF+POA****

stset spell_end_date, failure(cause_endpoint_enc==2 4) exit(failure) origin(time monit_origin_date) time0(spell_start_date) id(wolf_ID)

*Model 1 -- only treatment (lib_kill)

stcox i.lib_kill, efron robust cluster(wolf_ID)

estat ic

*checking PH assumption of treatment

stcox i.lib_kill, tvc(i.lib_kill) efron robust cluster(wolf_ID)

estat ic

*adding covariates*

*Model 2

stcox i.lib_kill i.winter, efron robust cluster(wolf_ID)

estat ic

*Model 3

stcox i.lib_kill i.winter, tvc(i.lib_kill i.winter) efron robust cluster(wolf_ID)

estat ic

*Model 4

stcox i.lib_kill i.sex_coded, efron robust cluster(wolf_ID)

estat ic

*Model 5

stcox i.lib_kill i.sex_coded, tvc(i.lib_kill i.sex_coded) efron robust cluster(wolf_ID)

estat ic

*BEST MODEL (#1)

stcox lib_kill, efron robust cluster(wolf_ID)

estat ic

stcurve, cumh at1(lib_kill=0) at2(lib_kill=1) legend(label(1 "Full Protections") label(2 "Reduced protections")) lpattern(solid dash) lcolor(navy maroon)

graph save "STCOX_CUMHAZ_LTFPOA", replace

*Checking PH assumptions -- TO USE WITH BEST MODEL (with that model run immediately preceding it)

estat phtest, log detail

estat phtest, plot(lib_kill)

graph save "STCOX_phtest_LTFPOA", replace

stphplot, by(lib_kill) nolnt

*Goodness of fit -- TO USE WITH BEST MODEL (look for NA line following Cox-snell residual line ~45 degree angle; from https://stats.idre.ucla.edu/stata/seminars/stata-survival)

quietly stcox lib_kill, nohr efron robust cluster(wolf_ID) mgale(mg)

predict cs, csnell

stset cs, id(wolf_ID) failure(cause_endpoint_enc==3)

sts generate H = na

line H cs cs, sort xlab(0 1 to 4) ylab(0 1 to 4)

drop mg cs H

graph save "STCOX_Cox-Snell_LTF", replace

*predicting HR for CIFs later -- TO USE WITH BEST MODEL

stset spell_end_date, failure(cause_endpoint_enc==2 4) exit(failure) origin(time monit_origin_date) time0(spell_start_date) id(wolf_ID)

stcox lib_kill, nohr efron robust cluster(wolf_ID)

predict h_ltfpoa_0, basehc

gsort _t -_d

by _t: replace h_ltfpoa_0 = . if _n > 1

gen h_ltfpoa_1 = h_ltfpoa_0*exp(_b[lib_kill])

twoway line h_ltfpoa_* _t, connect(J J) sort lpattern(solid dash) lcolor(navy maroon) ///

legend(label(1 "Full Protections") label(2 "Reduced protections")) ///

graphregion(color(white)) title("LTF Hazards STCOX")

graph save "STCOX_Hazards_LTFPOA", replace

**LTF**

stset spell_end_date, failure(cause_endpoint_enc==2) exit(failure) origin(time monit_origin_date) time0(spell_start_date) id(wolf_ID)

stset _t, failure(cause_endpoint_enc==2) exit(failure) origin(time _t0) time0(_t0) id(wolf_ID)

*Model 1 -- only treatment (lib_kill)

stcox i.lib_kill, nohr efron robust cluster(wolf_ID)

estat ic

*checking PH assumption of treatment

stcox i.lib_kill, tvc(i.lib_kill) efron robust cluster(wolf_ID)

estat ic

*adding covariates*

*Model 2

stcox i.lib_kill i.winter, efron robust cluster(wolf_ID)

estat ic

*Model 3

stcox i.lib_kill i.winter, tvc(i.lib_kill i.winter) efron robust cluster(wolf_ID)

estat ic

*Model 4

stcox i.lib_kill i.sex_coded, efron robust cluster(wolf_ID)

estat ic

*Model 5

stcox i.lib_kill i.sex_coded, tvc(i.lib_kill i.sex_coded) efron robust cluster(wolf_ID)

estat ic

*BEST MODEL (#1)

stcox lib_kill, efron robust cluster(wolf_ID)

estat ic

stcurve, cumh at1(lib_kill=0) at2(lib_kill=1)

graph save "STCOX_CUMHAZ_LTF", replace

*Checking PH assumptions -- TO USE WITH BEST MODEL (with that model run immediately preceding it)

estat phtest, log detail

estat phtest, plot(lib_kill)

graph save "STCOX_phtest_LTF", replace

stphplot, by(lib_kill) nolnt

*Goodness of fit -- TO USE WITH BEST MODEL (look for NA line following Cox-snell residual line ~45 degree angle; from https://stats.idre.ucla.edu/stata/seminars/stata-survival)

quietly stcox lib_kill, nohr efron robust cluster(wolf_ID) mgale(mg)

predict cs, csnell

stset cs, id(wolf_ID) failure(cause_endpoint_enc==2)

sts generate H = na

line H cs cs, sort xlab(0 1 to 2.5) ylab(0 1 to 2.5)

drop mg cs H

graph save "STCOX_Cox-Snell_LTF", replace

*predicting HR for CIFs later -- TO USE WITH BEST MODEL

stset spell_end_date, failure(cause_endpoint_enc==2) exit(failure) origin(time monit_origin_date) time0(spell_start_date) id(wolf_ID)

stcox lib_kill, nohr efron robust cluster(wolf_ID)

predict h_ltf_0, basehc

gsort _t -_d

by _t: replace h_ltf_0 = . if _n > 1

gen h_ltf_1 = h_ltf_0*exp(_b[lib_kill])

twoway line h_ltf_* _t, connect(J J) sort lpattern(solid dash) lcolor(navy maroon) ///

legend(label(1 "Full Protections") label(2 "Reduced protections")) ///

graphregion(color(white)) title("LTF Hazards STCOX")

graph save "STCOX_Hazards_LTF", replace

**ILLEGAL TAKE**

stset spell_end_date, failure(cause_endpoint_enc==4) exit(failure) origin(time monit_origin_date) time0(spell_start_date) id(wolf_ID)

*Model 1 -- only treatment (lib_kill)

stcox i.lib_kill, efron robust cluster(wolf_ID)

estat ic

*checking PH assumption of treatment

stcox i.lib_kill, tvc(i.lib_kill) efron robust cluster(wolf_ID)

estat ic

*adding covariates*

*Model 2

stcox i.lib_kill i.winter, efron robust cluster(wolf_ID)

estat ic

*Model 3

stcox i.lib_kill i.winter, tvc(i.lib_kill i.winter) efron robust cluster(wolf_ID)

estat ic

*Model 4

stcox i.lib_kill i.sex_coded, efron robust cluster(wolf_ID)

estat ic

*Model 5

stcox i.lib_kill i.sex_coded, tvc(i.lib_kill i.sex_coded) efron robust cluster(wolf_ID)

estat ic

*BEST MODEL (#1)

stcox lib_kill, efron robust cluster(wolf_ID)

estat ic

stcurve, cumh at1(lib_kill=0) at2(lib_kill=1)

graph save "STCOX_CUMHAZ_POA", replace

*Checking PH assumptions -- TO USE WITH BEST MODEL (with that model run immediately preceding it)

estat phtest, log detail

estat phtest, plot(lib_kill)

graph save "STCOX_phtest_POA", replace

stphplot, by(lib_kill) nolnt

*Goodness of fit -- TO USE WITH BEST MODEL (look for NA line following Cox-snell residual line ~45 degree angle)

quietly stcox lib_kill, nohr efron robust cluster(wolf_ID) mgale(mg)

predict cs, csnell

stset cs, id(wolf_ID) failure(cause_endpoint_enc==4)

sts generate H = na

line H cs cs, sort xlab(0 1 to 2.5) ylab(0 1 to 2.5)

drop mg cs H

graph save "STCOX_Cox-Snell_POA", replace

*predicting HR for CIFs later -- TO USE WITH BEST MODEL

stset spell_end_date, failure(cause_endpoint_enc==4) exit(failure) origin(time monit_origin_date) time0(spell_start_date) id(wolf_ID)

stcox lib_kill, nohr efron robust cluster(wolf_ID)

predict h_poa_0, basehc

gsort _t -_d

by _t: replace h_poa_0 = . if _n > 1

gen h_poa_1 = h_poa_0*exp(_b[lib_kill])

twoway line h_poa_* _t, connect(J J) sort lpattern(solid dash) lcolor(navy maroon) ///

legend(label(1 "Full Protections") label(2 "Reduced protections")) ///

graphregion(color(white)) title("Poached Hazards STCOX")

graph save "STCOX_Hazards_POA", replace

**AGENCY REMOVAL**

stset spell_end_date, failure(cause_endpoint_enc==3) exit(failure) origin(time monit_origin_date) time0(spell_start_date) id(wolf_ID)

*Model 1 -- only treatment (lib_kill)

stcox i.lib_kill, efron robust cluster(wolf_ID)

estat ic

*checking PH assumption of treatment

stcox i.lib_kill, tvc(i.lib_kill) efron robust cluster(wolf_ID)

estat ic

*adding covariates*

*Model 2

stcox i.lib_kill i.winter, efron robust cluster(wolf_ID)

estat ic

*Model 3

stcox i.lib_kill i.winter, tvc(i.lib_kill i.winter) efron robust cluster(wolf_ID)

estat ic

*Model 4

stcox i.lib_kill i.sex_coded, efron robust cluster(wolf_ID)

estat ic

*Model 5

stcox i.lib_kill i.sex_coded, tvc(i.lib_kill i.sex_coded) efron robust cluster(wolf_ID)

estat ic

*BEST MODEL (#1)

stcox lib_kill, efron robust cluster(wolf_ID)

estat ic

stcurve, cumh at1(lib_kill=0) at2(lib_kill=1)

graph save "STCOX_CUMHAZ_AR", replace

*Checking PH assumptions -- TO USE WITH BEST MODEL (with that model run immediately preceding it)

estat phtest, log detail

estat phtest, plot(lib_kill)

graph save "STCOX_phtest_AR", replace

stphplot, by(lib_kill) nolnt

*Goodness of fit -- TO USE WITH BEST MODEL (look for NA line following Cox-snell residual line ~45 degree angle)

quietly stcox lib_kill, nohr efron robust cluster(wolf_ID) mgale(mg)

predict cs, csnell

stset cs, id(wolf_ID) failure(cause_endpoint_enc==3)

sts generate H = na

line H cs cs, sort xlab(0 1 to 2.5) ylab(0 1 to 2.5)

drop mg cs H

graph save "STCOX_Cox-Snell_AR", replace

*predicting HR for CIFs later -- TO USE WITH BEST MODEL

stset spell_end_date, failure(cause_endpoint_enc==3) exit(failure) origin(time monit_origin_date) time0(spell_start_date) id(wolf_ID)

stcox lib_kill, nohr efron robust cluster(wolf_ID)

predict h_ar_0, basehc

gsort _t -_d

by _t: replace h_ar_0 = . if _n > 1

gen h_ar_1 = h_ar_0*exp(_b[lib_kill])

twoway line h_ar_* _t, connect(J J) sort lpattern(solid dash) lcolor(navy maroon) ///

legend(label(1 "Full Protections") label(2 "Reduced protections")) ///

graphregion(color(white)) title("Agency removal Hazards STCOX")

graph save "STCOX_Hazards_AR", replace

**NATURAL**

stset spell_end_date, failure(cause_endpoint_enc==5) exit(failure) origin(time monit_origin_date) time0(spell_start_date) id(wolf_ID)

*Model 1 -- only treatment (lib_kill)

stcox i.lib_kill, efron robust cluster(wolf_ID)

estat ic

*checking PH assumption of treatment

stcox i.lib_kill, tvc(i.lib_kill) efron robust cluster(wolf_ID)

estat ic

*adding covariates*

*Model 2

stcox i.lib_kill i.winter, efron robust cluster(wolf_ID)

estat ic

*Model 3

stcox i.lib_kill i.winter, tvc(i.lib_kill i.winter) efron robust cluster(wolf_ID)

estat ic

*Model 4

stcox i.lib_kill i.sex_coded, efron robust cluster(wolf_ID)

estat ic

*Model 5

stcox i.lib_kill i.sex_coded, tvc(i.lib_kill i.sex_coded) efron robust cluster(wolf_ID)

estat ic

*BEST MODEL (#1)

stcox lib_kill, efron robust cluster(wolf_ID)

estat ic

stcurve, cumh at1(lib_kill=0) at2(lib_kill=1) lpattern(solid dash) lcolor(navy maroon) legend(label(1 "Full Protections") label(2 "Reduced protections"))

graph save "STCOX_CUMHAZ_NAT", replace

*Checking PH assumptions -- TO USE WITH BEST MODEL (with that model run immediately preceding it)

estat phtest, log detail

estat phtest, plot(lib_kill)

graph save "STCOX_phtest_NAT", replace

stphplot, by(lib_kill) nolnt

*Goodness of fit -- TO USE WITH BEST MODEL (look for NA line following Cox-snell residual line ~45 degree angle)

quietly stcox lib_kill, nohr efron robust cluster(wolf_ID) mgale(mg)

predict cs, csnell

stset cs, id(wolf_ID) failure(cause_endpoint_enc==5)

sts generate H = na

line H cs cs, sort xlab(0 1 to 2.5) ylab(0 1 to 2.5)

drop mg cs H

graph save "STCOX_Cox-Snell_NAT", replace

*predicting HR for CIFs later -- TO USE WITH BEST MODEL

stset spell_end_date, failure(cause_endpoint_enc==5) exit(failure) origin(time monit_origin_date) time0(spell_start_date) id(wolf_ID)

stcox lib_kill, nohr efron robust cluster(wolf_ID)

predict h_nat_0, basehc

gsort _t -_d

by _t: replace h_nat_0 = . if _n > 1

gen h_nat_1 = h_nat_0*exp(_b[lib_kill])

twoway line h_nat_* _t, connect(J J) sort lpattern(solid dash) lcolor(navy maroon) ///

legend(label(1 "Full Protections") label(2 "Reduced protections")) ///

graphregion(color(white)) title("Natural Hazards STCOX")

graph save "STCOX_Hazards_NAT", replace

**NON-CRIMINAL**

stset spell_end_date, failure(cause_endpoint_enc==6) exit(failure) origin(time monit_origin_date) time0(spell_start_date) id(wolf_ID)

*Model 1 -- only treatment (lib_kill)

stcox i.lib_kill, efron robust cluster(wolf_ID)

estat ic

*checking PH assumption of treatment

stcox i.lib_kill, tvc(i.lib_kill) efron robust cluster(wolf_ID)

estat ic

*adding covariates*

*Model 2

stcox i.lib_kill i.winter, efron robust cluster(wolf_ID)

estat ic

*Model 3

stcox i.lib_kill i.winter, tvc(i.lib_kill i.winter) efron robust cluster(wolf_ID)

estat ic

*Model 4

stcox i.lib_kill i.sex_coded, efron robust cluster(wolf_ID)

estat ic

*Model 5

stcox i.lib_kill i.sex_coded, tvc(i.lib_kill i.sex_coded) efron robust cluster(wolf_ID)

estat ic

*BEST MODEL (#1)

stcox lib_kill, efron robust cluster(wolf_ID)

estat ic

stcurve, cumh at1(lib_kill=0) at2(lib_kill=1) lpattern(solid dash) lcolor(navy maroon) legend(label(1 "Full Protections") label(2 "Reduced protections"))

graph save "STCOX_CUMHAZ_NON", replace

*Checking PH assumptions -- TO USE WITH BEST MODEL (with that model run immediately preceding it)

estat phtest, log detail

estat phtest, plot(lib_kill)

graph save "STCOX_phtest_NON", replace

stphplot, by(lib_kill) nolnt

*Goodness of fit -- TO USE WITH BEST MODEL (look for NA line following Cox-snell residual line ~45 degree angle)

quietly stcox lib_kill, nohr efron robust cluster(wolf_ID) mgale(mg)

predict cs, csnell

stset cs, id(wolf_ID) failure(cause_endpoint_enc==6)

sts generate H = na

line H cs cs, sort xlab(0 1 to 2.5) ylab(0 1 to 2.5)

drop mg cs H

graph save "STCOX_Cox-Snell_NON", replace

*predicting HR for CIFs later -- TO USE WITH BEST MODEL

stset spell_end_date, failure(cause_endpoint_enc==6) exit(failure) origin(time monit_origin_date) time0(spell_start_date) id(wolf_ID)

stcox lib_kill, nohr efron robust cluster(wolf_ID)

predict h_non_0, basehc

gsort _t -_d

by _t: replace h_non_0 = . if _n > 1

gen h_non_1 = h_non_0*exp(_b[lib_kill])

twoway line h_non_* _t, connect(J J) sort lpattern(solid dash) lcolor(navy maroon) ///

legend(label(1 "Full Protections") label(2 "Reduced protections")) ///

graphregion(color(white)) title("Non-criminal Hazards STCOX")

graph save "STCOX_Hazards_NON", replace

********************************************************************************

**CREATING CIFs WITH ABOVE HAZARD CONTRIBUTIONS (see competingrisk_statuse.pdf)*

drop if missing(h_ltf_0) & missing(h_ar_0) & missing(h_poa_0) & missing(h_nat_0) & missing(h_non_0)

replace h_ltf_0=0 if missing(h_ltf_0)

replace h_ltf_1=0 if missing(h_ltf_1)

replace h_ar_0=0 if missing(h_ar_0)

replace h_ar_1=0 if missing(h_ar_1)

replace h_poa_0=0 if missing(h_poa_0)

replace h_poa_1=0 if missing(h_poa_1)

replace h_nat_0=0 if missing(h_nat_0)

replace h_nat_1=0 if missing(h_nat_1)

replace h_non_0=0 if missing(h_non_0)

replace h_non_1=0 if missing(h_non_1)

**calculating event-free survivor functions

sort _t

gen S_0 = exp(sum(log(1- h_ltf_0 – h_poa_0 - h_ar_0 - h_nat_0 - h_non_0)))

gen S_1 = exp(sum(log(1- h_ltf_1 – h_poa_1 - h_ar_1 - h_nat_1 - h_non_1)))

twoway line S_* _t, connect(J J) sort

*calculating CIFs

*LTF

gen cif_ltf_0 = sum(S_0[_n-1]*h_ltf_0)

gen cif_ltf_1 = sum(S_1[_n-1]*h_ltf_1)

twoway line cif_ltf_* _t, connect(J J) sort lpattern(solid dash) lcolor(navy maroon) ///

legend(label(1 "Full Protections") label(2 "Reduced protections")) ///

graphregion(color(white)) title("LTF CIFs STCOX")

graph save "STCOX_CIFs_LTF", replace

*Illegal take

gen cif_poa_0 = sum(S_0[_n-1]*h_poa_0)

gen cif_poa_1 = sum(S_1[_n-1]*h_poa_1)

twoway line cif_poa_* _t, connect(J J) sort lpattern(solid dash) lcolor(navy maroon) ///

legend(label(1 "Full Protections") label(2 "Reduced protections")) ///

graphregion(color(white)) ytitle("Cumulative Incidence") title("POA CIFs STCOX")

graph save "STCOX_CIFs_POA", replace

*Agency removal

gen cif_ar_0 = sum(S_0[_n-1]*h_ar_0)

gen cif_ar_1 = sum(S_1[_n-1]*h_ar_1)

twoway line cif_ar_* _t, connect(J J) sort lpattern(solid dash) lcolor(navy maroon) ///

legend(label(1 "Full Protections") label(2 "Reduced protections")) ///

graphregion(color(white)) ytitle("Cumulative Incidence") title("AR CIFs STCOX")

graph save "STCOX_CIFs_AR", replace

*Natural

gen cif_nat_0 = sum(S_0[_n-1]*h_nat_0)

gen cif_nat_1 = sum(S_1[_n-1]*h_nat_1)

twoway line cif_nat_* _t, connect(J J) sort lpattern(solid dash) lcolor(navy maroon) ///

legend(label(1 "Full Protections") label(2 "Reduced protections")) ///

graphregion(color(white)) ytitle("Cumulative Incidence") title("Natural CIFs STCOX")

graph save "STCOX_CIFs_NAT", replace

*Non-criminal

gen cif_non_0 = sum(S_0[_n-1]*h_non_0)

gen cif_non_1 = sum(S_1[_n-1]*h_non_1)

twoway line cif_non_* _t, connect(J J) sort lpattern(solid dash) lcolor(navy maroon) ///

legend(label(1 "Full Protections") label(2 "Reduced protections")) ///

graphregion(color(white)) ytitle("Cumulative Incidence") title("Non-criminal CIFs STCOX")

graph save "STCOX_CIFs_NON", replace

********************************************************************************

***SEMI-PARAMETRIC APPROACHES FOR CIFS******************************************

***FINE & GRAY APPROACH FOR COMPETING RISK, BY CAUSE OF FAILURE*****************

***LTF+POA***

stset spell_end_date, failure(cause_endpoint_enc==2 4) exit(failure) origin(time monit_origin_date) time0(spell_start_date) id(wolf_ID)

*Model 1

stcrreg i.lib_kill, compete(cause_endpoint_enc==3 5 6) nolog show vce(cluster wolf_ID)

estat ic

stcrreg i.lib_kill, tvc(i.lib_kill) compete(cause_endpoint_enc==2 4 5) nolog show vce(cluster wolf_ID)

estat ic

**adding covariates**

*Model 2

stcrreg i.lib_kill i.winter, compete(cause_endpoint_enc==2 4 5) nolog show vce(cluster wolf_ID)

estat ic

*Model 3

stcrreg i.lib_kill i.winter i.sex_coded, compete(cause_endpoint_enc==2 4 5) nolog show vce(cluster wolf_ID)

estat ic

*Model 4

stcrreg i.lib_kill i.sex_coded, compete(cause_endpoint_enc==2 4 5) nolog show vce(cluster wolf_ID)

estat ic

estat ic

*BEST MODEL (#1)

stcrreg i.lib_kill, compete(cause_endpoint_enc==3 5 6) nolog show vce(cluster wolf_ID)

estat ic

/*ANALYSIS --> */

*using best models to calculate CIFs*?

stcrreg i.lib_kill, compete(cause_endpoint_enc==3 5 6) nolog show vce(cluster wolf_ID)

stcurve, cif at1(lib_kill = 0) at2(lib_kill = 1) lpattern(solid dash) lcolor(navy maroon) ///

legend(label(1 "Full Protections") label(2 "Reduced protections")) ///

graphregion(color(white)) title("LTFPOA CIFs F&G")

graph save "F&G_CIFs_LTFPOA", replace

predict fg_ltfpoa_0, basecif

gsort _t -_d

by _t: replace fg_ltfpoa_0 = . if _n > 1

gen fg_ltfpoa_1 = fg_ltfpoa_0*exp(_b[1.lib_kill])

twoway line fg_ar_* fg_ltfpoa_* _t, connect(J J) sort lpattern(solid solid longdash longdash shortdash shortdash) lcolor(navy maroon navy maroon navy maroon) ///

legend(label(1 "AR w/Protections") label(2 "AR w/Killing") label(3 "LTFPOA w/Protections") label(4 "LTFPOA w/Killing")) ///

graphregion(color(white)) ytitle("Cumulative Incidence") title("AR,LTFPOA CIFs FG")

graph save "FG_CIFs_AR_LTFPOA_combined", replace

graph export "FG_CIFs_AR_LTFPOA_combined.pdf", replace

graph combine "F&G_CIFs_LTFPOA" "STCOX_CIFs_LTFPOA", com saving("F&G_STCOX_CIFs_LTFPOA")

graph export "F&G_STCOX_CIFs_LTFPOA.pdf", replace

**LTF**

stset spell_end_date, failure(cause_endpoint_enc==2) exit(failure) origin(time monit_origin_date) time0(spell_start_date) id(wolf_ID)

*Model 1

stcrreg i.lib_kill, compete(cause_endpoint_enc==3 4 5 6) nolog show vce(cluster wolf_ID)

estat ic

stcrreg i.lib_kill, tvc(i.lib_kill) compete(cause_endpoint_enc==3 4 5 6) nolog show vce(cluster wolf_ID)

estat ic

**adding covariates**

*Model 2

stcrreg i.lib_kill i.winter, compete(cause_endpoint_enc==3 4 5 6) nolog show vce(cluster wolf_ID)

estat ic

*Model 3

stcrreg i.lib_kill i.winter i.sex_coded, compete(cause_endpoint_enc==3 4 5 6) nolog show vce(cluster wolf_ID)

estat ic

*Model 4

stcrreg i.lib_kill i.winter i.sex_coded, tvc(i.lib_kill i.winter i.sex_coded) compete(cause_endpoint_enc==3 4 5 6) nolog show vce(cluster wolf_ID)

estat ic

*Model 5

stcrreg i.lib_kill i.sex_coded, compete(cause_endpoint_enc==3 4 5 6) nolog show vce(cluster wolf_ID)

estat ic

*BEST MODEL (#5)

stcrreg i.lib_kill i.sex_coded, compete(cause_endpoint_enc==3 4 5 6) nolog show vce(cluster wolf_ID)

estat ic

/*ANALYSIS --> */

*using best model to calculate CIFs

stcrreg i.lib_kill i.sex_coded, compete(cause_endpoint_enc==3 4 5 6) nolog show vce(cluster wolf_ID)

stcurve, cif at1(lib_kill = 0) at2(lib_kill = 1) lpattern(solid dash) lcolor(navy maroon) ///

legend(label(1 "Full Protections") label(2 "Reduced protections")) ///

graphregion(color(white)) title("LTF CIFs F&G")

graph save "F&G_CIFs_LTF", replace

predict fg_ltf_0, basecif

gsort _t -_d

by _t: replace fg_ltf_0 = . if _n > 1

gen fg_ltf_1 = fg_ltf_0*exp(_b[1.lib_kill])

**ILLEGAL TAKE**

stset spell_end_date, failure(cause_endpoint_enc==4) exit(failure) origin(time monit_origin_date) time0(spell_start_date) id(wolf_ID)

*Model 1

stcrreg i.lib_kill, compete(cause_endpoint_enc==2 3 5 6) nolog show vce(cluster wolf_ID)

estat ic

stcrreg i.lib_kill, tvc(i.lib_kill) compete(cause_endpoint_enc==2 3 5 6) nolog show vce(cluster wolf_ID)

estat ic

**adding covariates**

*Model 2

stcrreg i.lib_kill i.winter, compete(cause_endpoint_enc==2 3 5 6) nolog show vce(cluster wolf_ID)

estat ic

*Model 3

stcrreg i.lib_kill i.winter i.sex_coded, compete(cause_endpoint_enc==2 3 5 6) nolog show vce(cluster wolf_ID)

estat ic

*Model 4

stcrreg i.lib_kill i.sex_coded, compete(cause_endpoint_enc==2 3 5 6) nolog show vce(cluster wolf_ID)

estat ic

*estat ic

*BEST MODEL (#1)

stcrreg i.lib_kill, compete(cause_endpoint_enc==2 3 5 6) nolog show vce(cluster wolf_ID)

estat ic

/*ANALYSIS --> */

*using best models to calculate CIFs*?

stcrreg i.lib_kill, compete(cause_endpoint_enc==2 3 5 6) nolog show vce(cluster wolf_ID)

stcurve, cif at1(lib_kill = 0) at2(lib_kill = 1) lpattern(solid dash) lcolor(navy maroon) ///

legend(label(1 "Full Protections") label(2 "Reduced protections")) ///

graphregion(color(white)) title("POA CIFs F&G")

*graph save "F&G_CIFs_POA", replace

predict fg_poa_0, basecif

gsort _t -_d

by _t: replace fg_poa_0 = . if _n > 1

gen fg_poa_1 = fg_poa_0*exp(_b[1.lib_kill])

twoway line fg_ar_* fg_ltf_* fg_poa_* _t, connect(J J) sort lpattern(solid solid longdash longdash shortdash shortdash) lcolor(navy maroon navy maroon navy maroon) ///

legend(label(1 "AR w/Protections") label(2 "AR w/Killing") label(3 "LTF w/Protections") label(4 "LTF w/Killing") label(5 "Poached w/Protections") label(6 "Poached w/Killing")) ///

graphregion(color(white)) ytitle("Cumulative Incidence") title("AR,LTF,POA CIFs FG")

*graph save "FG_CIFs_AR_LTF_POA_combined", replace

*graph export "FG_CIFs_AR_LTF_POA_combined.pdf", replace

graph combine "F&G_CIFs_POA" "STCOX_CIFs_POA", com saving("F&G_STCOX_CIFs_POA")

graph export "F&G_STCOX_CIFs_POA.pdf", replace

/*ADJUST FOR FINAL GRAPH*

twoway line fg_ar_* fg_ltf_* fg_poa_* _t , connect(J J) sort lpattern(solid dash solid dash solid dash) lcolor(black black dkorange dkorange maroon maroon) ///

legend(off) graphregion(color(white)) ytitle("Cumulative Incidence") xtitle("time when monitoring ends (days)") text(.45 2500 "LTF", place(ne) color(dkorange)) text(.0 2500 "AR", place(ne) color(black)) ///

text(.05 2500 "Poached", place(ne) color(maroon))*/

**AGENCY REMOVAL**

stset spell_end_date, failure(cause_endpoint_enc==3) exit(failure) origin(time monit_origin_date) time0(spell_start_date) id(wolf_ID)

*Model 1

stcrreg i.lib_kill, compete(cause_endpoint_enc==2 4 5 6) nolog show vce(cluster wolf_ID)

estat ic

stcrreg i.lib_kill, tvc(i.lib_kill) compete(cause_endpoint_enc==2 4 5 6) nolog show vce(cluster wolf_ID)

estat ic

**adding covariates**

*Model 2

stcrreg i.lib_kill i.winter, compete(cause_endpoint_enc==2 4 5 6) nolog show vce(cluster wolf_ID)

estat ic

*Model 3

stcrreg i.lib_kill i.winter i.sex_coded, compete(cause_endpoint_enc==2 4 5 6) nolog show vce(cluster wolf_ID)

estat ic

*Model 4

stcrreg i.lib_kill i.sex_coded, compete(cause_endpoint_enc==2 4 5 6) nolog show vce(cluster wolf_ID)

estat ic

*BEST MODEL (#1)

stcrreg i.lib_kill, compete(cause_endpoint_enc==2 4 5 6) nolog show vce(cluster wolf_ID)

estat ic

/*ANALYSIS --> */

*using best model to calculate CIFs

stcrreg i.lib_kill, compete(cause_endpoint_enc==2 4 5 6) nolog show vce(cluster wolf_ID)

stcurve, cif at1(lib_kill = 0) at2(lib_kill = 1) lpattern(solid dash) lcolor(navy maroon) ///

legend(label(1 "Full Protections") label(2 "Reduced protections")) ///

graphregion(color(white)) title("AR CIFs F&G")

graph save "F&G_CIFs_AR", replace

graph combine "F&G_CIFs_AR" "STCOX_CIFs_AR", com saving("F&G_STCOX_CIFs_AR")

graph export "F&G_STCOX_CIFs_AR.pdf", replace

graph combine "F&G_CIFs_AR" "F&G_CIFs_LTF", com saving("F&G_CIFs_LTF_AR")

graph export "F&G_CIFs_LTF_AR.pdf", replace

predict fg_ar_0, basecif

gsort _t -_d

by _t: replace fg_ar_0 = . if _n > 1

gen fg_ar_1 = fg_ar_0*exp(_b[1.lib_kill])

twoway line fg_ar_* fg_ltf_* _t, connect(J J) sort lpattern(solid solid longdash longdash) lcolor(navy maroon navy maroon) ///

legend(label(1 "AR w/Protections") label(2 "AR w/Killing") label(3 "LTFPOA w/Protections") label(4 "LTFPOA w/Killing")) ///

graphregion(color(white)) ytitle("Cumulative Incidence") title("AR-LTF CIFs FG")

graph save "FG_CIFs_AR_LTFPOA_combined", replace

graph export "FG_CIFs_AR_LTFPOA_combined.pdf", replace

**NATURAL**

stset spell_end_date, failure(cause_endpoint_enc==5) exit(failure) origin(time monit_origin_date) time0(spell_start_date) id(wolf_ID)

*Model 1

stcrreg i.lib_kill, compete(cause_endpoint_enc==2 3 4 6) nolog show vce(cluster wolf_ID)

estat ic

stcrreg i.lib_kill, tvc(i.lib_kill) compete(cause_endpoint_enc==2 3 4 6) nolog show vce(cluster wolf_ID)

estat ic

**adding covariates**

*Model 2

stcrreg i.lib_kill i.winter, compete(cause_endpoint_enc==2 3 4 6) nolog show vce(cluster wolf_ID)

estat ic

*Model 3

stcrreg i.lib_kill i.sex_coded, compete(cause_endpoint_enc==2 3 4 6) nolog show vce(cluster wolf_ID)

estat ic

*Model 4

stcrreg i.lib_kill i.winter i.sex_coded, tvc(i.lib_kill i.winter i.sex_coded) compete(cause_endpoint_enc==2 3 4 6) nolog show vce(cluster wolf_ID)

estat ic

*BEST MODEL (#1)

stcrreg i.lib_kill, compete(cause_endpoint_enc==2 3 4 6) nolog show vce(cluster wolf_ID)

estat ic

/*ANALYSIS --> */

*using best model to calculate CIFs

stcrreg i.lib_kill, compete(cause_endpoint_enc==2 3 4 6) nolog show vce(cluster wolf_ID)

stcurve, cif at1(lib_kill = 0) at2(lib_kill = 1) lpattern(solid dash) lcolor(navy maroon) ///

legend(label(1 "Full Protections") label(2 "Reduced protections")) ///

graphregion(color(white)) title("Natural CIFs F&G")

graph save "F&G_CIFs_NAT", replace

graph combine "F&G_CIFs_NAT" "STCOX_CIFs_NAT", com saving("F&G_STCOX_CIFs_NAT")

graph export "F&G_STCOX_CIFs_NAT.pdf", replace

predict fg_nat_0, basecif

gsort _t -_d

by _t: replace fg_nat_0 = . if _n > 1

gen fg_nat_1 = fg_nat_0*exp(_b[1.lib_kill])

twoway line fg_ar_* fg_ltf_* fg_poa_* fg_nat_* _t , connect(J J) sort lpattern(solid solid longdash longdash shortdash shortdash dash_dot dash_dot) lcolor(navy maroon navy maroon navy maroon navy maroon) ///

legend(label(1 "AR w/Protections") label(2 "AR w/Killing") label(3 "LTF w/Protections") label(4 "LTF w/Killing") label(5 "Poached w/Protections") label(6 "Poached w/Killing") ///

label(7 "Natural w/Protections") label(8 "Natural w/Killing")) graphregion(color(white)) ytitle("Cumulative Incidence") title("AR,LTF,POA,NAT CIFs FG")

graph save "CIFs_AR_LTF_POA_NAT_combined", replace

graph export "CIFs_AR_LTF_POA_NAT_combined.pdf", replace

/*ADJUST FOR FINAL GRAPH*

twoway line fg_ar_* fg_ltf_* fg_poa_* fg_nat_* _t , connect(J J) sort lpattern(solid solid longdash longdash shortdash shortdash dash_dot dash_dot) lcolor(navy maroon navy maroon navy maroon navy maroon) ///

legend(label(1 "AR w/Protections") label(2 "AR w/Killing") label(3 "LTF w/Protections") label(4 "LTF w/Killing") label(5 "Poached w/Protections") label(6 "Poached w/Killing") ///

label(7 "Natural w/Protections") label(8 "Natural w/Killing") rows(2) cols(4) size(small)) graphregion(color(white)) ytitle("Cumulative Incidence")*/

**NON-CRIMINAL**

stset spell_end_date, failure(cause_endpoint_enc==6) exit(failure) origin(time monit_origin_date) time0(spell_start_date) id(wolf_ID)

*Model 1

stcrreg i.lib_kill, compete(cause_endpoint_enc==2 3 4 5) nolog show vce(cluster wolf_ID)

estat ic

stcrreg i.lib_kill, tvc(i.lib_kill) compete(cause_endpoint_enc==2 3 4 5) nolog show vce(cluster wolf_ID)

estat ic

**adding covariates**

*Model 2

stcrreg i.lib_kill i.winter, compete(cause_endpoint_enc==2 3 4 5) nolog show vce(cluster wolf_ID)

estat ic

*Model 3

stcrreg i.lib_kill i.sex_coded, compete(cause_endpoint_enc==2 3 4 5) nolog show vce(cluster wolf_ID)

estat ic

*Model 4

stcrreg i.lib_kill i.winter i.sex_coded, tvc(i.lib_kill i.winter i.sex_coded) compete(cause_endpoint_enc==2 3 4 5) nolog show vce(cluster wolf_ID)

estat ic

*BEST MODEL (#1)

stcrreg i.lib_kill, compete(cause_endpoint_enc==2 3 4 5) nolog show vce(cluster wolf_ID)

estat ic

/*ANALYSIS --> */

*using best model to calculate CIFs --> *different model than STCOX (this one includes winter)*

stcrreg i.lib_kill, compete(cause_endpoint_enc==2 3 4 5) nolog show vce(cluster wolf_ID)

stcurve, cif at1(lib_kill = 0) at2(lib_kill = 1) lpattern(solid dash) lcolor(navy maroon) ///

legend(label(1 "Full Protections") label(2 "Reduced protections")) ///

graphregion(color(white)) title("Non-criminal CIFs F&G")

graph save "F&G_CIFs_NON", replace

graph combine "F&G_CIFs_NON" "STCOX_CIFs_NON", com saving("F&G_STCOX_CIFs_NON")

graph export "F&G_STCOX_CIFs_NON.pdf", replace

********************************************************************************

***CHECKING F&G/STCOX MODELS AGAINST NON-PARAMETRIC F&G MODELS******************

********************************************************************************

***LTF***

stset spell_end_date, failure(cause_endpoint_enc==2) exit(failure) origin(time monit_origin_date) time0(spell_start_date) id(wolf_ID)

stcrreg if lib_kill==0, compete(cause_endpoint_enc==3 4 5 6) nolog show vce(cluster wolf_ID)

predict nonp_cif_ltf_0, basecif

stcrreg if lib_kill==1, compete(cause_endpoint_enc==3 4 5 6) nolog show vce(cluster wolf_ID)

predict nonp_cif_ltf_1, basecif

twoway line nonp_cif_ltf_* _t, connect(J J) sort lpattern(solid dash) lcolor(navy maroon) ///

legend(label(1 "Full Protections") label(2 "Reduced protections")) ///

graphregion(color(white)) ytitle("Cumulative Incidence") title("LTF CIFs NONPAR")

graph save "NONPAR_CIFs_LTF", replace

graph combine "NONPAR_CIFs_LTF" "F&G_CIFs_LTF" "STCOX_CIFs_LTF", com saving("NONPAR_F&G_STCOX_CIFs_LTF")

graph export "NONPAR_F&G_STCOX_CIFs_LTF.pdf", replace

***AGENCY REMOVAL***

stset spell_end_date, failure(cause_endpoint_enc==3) exit(failure) origin(time monit_origin_date) time0(spell_start_date) id(wolf_ID)

stcrreg if lib_kill==0, compete(cause_endpoint_enc==2 4 5 6) nolog show vce(cluster wolf_ID)

predict nonp_cif_ar_0, basecif

stcrreg if lib_kill==1, compete(cause_endpoint_enc==2 4 5 6) nolog show vce(cluster wolf_ID)

predict nonp_cif_ar_1, basecif

twoway line nonp_cif_ar_* _t, connect(J J) sort lpattern(solid dash) lcolor(navy maroon) ///

legend(label(1 "Full Protections") label(2 "Reduced protections")) ///

graphregion(color(white)) ytitle("Cumulative Incidence") title("AR CIFs NONPAR")

graph save "NONPAR_CIFs_AR", replace

graph combine "NONPAR_CIFs_AR" "F&G_CIFs_AR" "STCOX_CIFs_AR", com saving("NONPAR_F&G_STCOX_CIFs_AR")

graph export "NONPAR_F&G_STCOX_CIFs_AR.pdf", replace

***ILLEGAL TAKE***

stset spell_end_date, failure(cause_endpoint_enc==4) exit(failure) origin(time monit_origin_date) time0(spell_start_date) id(wolf_ID)

stcrreg if lib_kill==0, compete(cause_endpoint_enc==2 3 5 6) nolog show vce(cluster wolf_ID)

predict nonp_cif_poa_0, basecif

stcrreg if lib_kill==1, compete(cause_endpoint_enc==2 3 5 6) nolog show vce(cluster wolf_ID)

predict nonp_cif_poa_1, basecif

twoway line nonp_cif_poa_* _t, connect(J J) sort lpattern(solid dash) lcolor(navy maroon) ///

legend(label(1 "Full Protections") label(2 "Reduced protections")) ///

graphregion(color(white)) ytitle("Cumulative Incidence") title("POA CIFs NONPAR")

graph save "NONPAR_CIFs_POA", replace

graph combine "NONPAR_CIFs_POA" "F&G_CIFs_POA" "STCOX_CIFs_POA", com saving("NONPAR_F&G_STCOX_CIFs_POA")

graph export "NONPAR_F&G_STCOX_CIFs_POA.pdf", replace

***NATURAL***

stset spell_end_date, failure(cause_endpoint_enc==5) exit(failure) origin(time monit_origin_date) time0(spell_start_date) id(wolf_ID)

stcrreg if lib_kill==0, compete(cause_endpoint_enc==2 3 4 6) nolog show vce(cluster wolf_ID)

predict nonp_cif_nat_0, basecif

stcrreg if lib_kill==1, compete(cause_endpoint_enc==2 3 4 6) nolog show vce(cluster wolf_ID)

predict nonp_cif_nat_1, basecif

twoway line nonp_cif_nat_* _t, connect(J J) sort lpattern(solid dash) lcolor(navy maroon) ///

legend(label(1 "Full Protections") label(2 "Reduced protections")) ///

graphregion(color(white)) ytitle("Cumulative Incidence") title("Natural CIFs NONPAR")

graph save "NONPAR_CIFs_NAT", replace

graph combine "NONPAR_CIFs_NAT" "F&G_CIFs_NAT" "STCOX_CIFs_NAT", com saving("NONPAR_F&G_STCOX_CIFs_NAT")

graph export "NONPAR_F&G_STCOX_CIFs_NAT.pdf", replace

***NON-CRIMINAL***

stset spell_end_date, failure(cause_endpoint_enc==6) exit(failure) origin(time monit_origin_date) time0(spell_start_date) id(wolf_ID)

stcrreg if lib_kill==0, compete(cause_endpoint_enc==2 3 4 5) nolog show vce(cluster wolf_ID)

predict nonp_cif_non_0, basecif

stcrreg if lib_kill==1, compete(cause_endpoint_enc==2 3 4 5) nolog show vce(cluster wolf_ID)

predict nonp_cif_non_1, basecif

twoway line nonp_cif_non_* _t, connect(J J) sort lpattern(solid dash) lcolor(navy maroon) ///

legend(label(1 "Full Protections") label(2 "Reduced protections")) ///

graphregion(color(white)) ytitle("Cumulative Incidence") title("Non-criminal CIFs NONPAR")

graph save "NONPAR_CIFs_NON", replace

graph combine "NONPAR_CIFs_NON" "F&G_CIFs_NON" "STCOX_CIFs_NON", com saving("NONPAR_F&G_STCOX_CIFs_NON")

graph export "NONPAR_F&G_STCOX_CIFs_NON.pdf", replace

log close
